# Longitudinal effects of antibiotics and fecal transplant on lemur gut microbiota structure, associations, and resistomes

**DOI:** 10.1101/2020.11.11.378349

**Authors:** Sally L. Bornbusch, Rachel L. Harris, Nicholas M. Grebe, Kimberly Roche, Kristin Dimac-Stohl, Christine M. Drea

## Abstract

Antibiotics alter the diversity, structure, and dynamics of host-associated microbial consortia, including via development of antibiotic resistance; however, patterns of recovery from dysbiosis and methods to mitigate negative effects, remain poorly understood. We applied an ecological framework via long-term, integrated study of community structure, across scales, to improve understanding of host-microbe symbiosis during dysbiosis and recovery. We experimentally administered a broad-spectrum antibiotic alone or with subsequent fecal transfaunation to healthy, male ring-tailed lemurs (*Lemur catta*) and longitudinally tracked the diversity, composition, associations, and resistomes of their gut microbiota. Whereas microbial diversity recovered rapidly in lemurs, antibiotics caused long-term instability in community composition – effects that were attenuated by fecal transfaunation. Antibiotic resistance genes, which were universally present, including in treatment-naïve subjects, increased during and persisted after antibiotic treatment. Long-term, integrated study post antibiotic-induced dysbiosis revealed differential, metric-dependent evidence of recovery, beneficial effects of fecal transfaunation, and negative consequences to lemur resistomes.

## Introduction

The long, co-evolutionary history between vertebrates and their microbes underpins the complex web of interactions linking commensal microbiota to host function^1,2^. Because perturbations to these communities can have both short- and long-term, negative consequences^3–5^, we increasingly recognize the benefits provided by our endogenous microbiota and have come to view them as ‘old friends’^6,7^. To exemplify, while antibiotic treatment effectively combats immediate bacterial infections, it can also lead to prolonged, severe negative side-effects, such as the elimination of beneficial microbes and the deterioration of microbiome function^8,9^. Moreover, antibiotics also promote changes in microbial genomes; the ubiquitous use of antibiotics has spurred the spread of genes encoding antibiotic resistance (ABR), which can have potentially catastrophic consequences^10^. Microbial therapies, such as fecal transfaunation, can mitigate the detrimental side-effects of antibiotics^11^; however, because antibiotics are often studied in the context of preexisting illness or injury (which independently influences microbial communities), the severity, duration, and recovery from dysbiosis owing purely to antibiotics remain unclear. Here, we apply an ecological framework in healthy animals to better understand the trajectory and processes governing recovery or return to a stable microbial community following antibiotic-induced dysbiosis. Because nonhuman primates are prime models in which to probe microbial dynamics and the development of ABR in response to antibiotic treatment, we experimentally administered a broad-spectrum antibiotic to male ring-tailed lemurs (*Lemur catta*) and used a longitudinal approach to track impacts on the composition and resistomes of their gut microbiota. We further tested the effects of fecal transfaunation as an intervention to promote the recovery of microbial composition and to potentially mitigate the development and persistence of ABR.

Antibiotics and ABR genes have ancient origins as natural compounds or genetic defenses, respectively, used by microbes to compete and survive in densely populated communities, whether within or outside of a host^12,13^. The ability of bacteria to rapidly undergo mutation^14,15^ and share advantageous genes via lateral gene transfer^16,17^ has resulted in myriad, naturally occurring ABR genes^18,19^. The response of a microbial community to natural antibiotics is largely dictated by the interactions between microbial taxa, which vary over time and across environments. The efficacy and ubiquity of man-made antibiotics have severely perturbed microbial communities via targeted (e.g. narrow spectrum) or indiscriminate (e.g. broad spectrum) elimination of bacterial groups^20^, thereby altering the composition and, ultimately, functional potential of microbiomes^20–22^. In addition, these antibiotics have magnified selective pressure on bacterial communities, making ABR genes advantageous and instigating their proliferation ^23,24^, thereby altering the microbiota’s genomic make-up. Within host-associated microbiomes, the propagation of ABR can result in virulent, resistant pathogens ^25,26^ that reduce the diversity of native or beneficial microbes^27,28^ and diminish immune capacity of the host. Our understanding of these phenomena primarily derives from studies that characterize the effects of antibiotics on the elimination or development of ABR within specific bacterial pathogens^29,30^. We know comparatively less about how man-made antibiotics influence the aggregate interactions within presumed healthy, host-associated communities and how those dynamics influence the recovery of microbiota.

Recognizing that commensal consortia are vital to the host has spurred increased research into microbial therapies to mitigate the negative consequences of dysbiosis. In fecal transfaunation, for example, a ‘healthy’ or ‘native’ community of microbes sourced from feces is transferred into a dysbiotic community to combat pathogens and promote the growth of beneficial microbes^31,32^. Because coprophagy (the ingestion of fecal material either directly or via prey consumption) bolsters gut microbiota during development or illness^33,34^, medical practitioners have examined the use of fecal transfaunations to treat gastrointestinal distress in a wide range of host taxa^35,36^. As in studies of antibiotics, however, the effects of fecal transfaunation are best understood in the context of infection (with e.g. *Clostridium difficile*^37,38^). Whether or not fecal transfaunation alters the trajectory of microbiome recovery more broadly remains unclear.

Understudied compared to anthropoid primates, lemurs have a unique evolutionary trajectory that makes them interesting models in which to study the dynamics between hosts and their co-evolved microbes^39–42^. Ring-tailed lemurs are ecologically flexible^43,44^, owing in part to their highly omnivorous diet, making them one of the few lemur species to thrive in captivity. This flexibility is reflected in their resilient gut microbiota that seem relatively unperturbed by aspects of captivity^42^. Ring-tailed lemurs re also relatively robust to health concerns, such as gastrointestinal problems, that affect the microbiota and welfare of other captive strepsirrhines^45^.

Here, we apply classic ecological principles to gut microbial communities to investigate two non-exclusive hypotheses regarding post-dysbiosis recovery. We use experimental manipulations (antibiotic treatment with or without fecal transfaunation), paired with longitudinal data to examine patterns in microbiota structure (e.g. alpha and beta diversity via 16S rRNA amplicon sequencing), bacterial associations (via Bayesian models of covariation), and ABR gene profiles (via shotgun metagenomic sequencing). First, diversity increases functional redundancy within a community and thus improves stability^46–48^. Under the ‘diversity begets stability’ hypothesis, as applied to a dysbiotic microbiome, recovery of alpha diversity, regardless of microbial identity, should be vital and sufficient to achieve a stable microbiome^49–51^. Accordingly, after antibiotic treatment, we would expect to see an increase in microbial richness (e.g. alpha diversity), independent of fecal transfaunation. The resulting stable communities of the two treatment groups could thus have similar richness, but different compositions. Alternately, previous evidence also indicates that certain community members (i.e., keystone species or specific ‘old friends’) are foundational to community function^52–55^, such that recovery of a stable microbiome requires specific community composition (e.g. beta diversity). Under the ‘key-stone species’ hypothesis, we would predict that, following antibiotic-mediated dysbiosis, there would be recovery of the same community composition, with fecal transfaunation accelerating the recovery rate. Accordingly, the resulting stable communities of both treatment groups would have similar compositions. These two hypotheses could also work in concert, but along different schedules, with potentially more rapid recovery of richness, but slower and more variable recovery of composition. Notably, the complexity of dynamics between specific community members (i.e., cooperation and competition) could create long-term fluctuations in community composition that would be highlighted by bacterial covariations between key members of the community. Furthermore, the presence of ABR within the microbiomes could exert a distinct force in driving community composition during the treatment and recovery phases. By tracking ABR prevalence and type, coupled with bacterial covariation, we can make inferences about which microbes may be harboring and expressing ABR genes.

## Results

### Baseline and control bacterial communities

In the pretreatment phase, neither alpha nor beta diversity varied significantly between the three experimental groups (alpha diversity: Kruskal-Wallis test, H = 2.478, p = 0.289, beta diversity: H = 2.658, p = 0.264). Across all control samples (across all phases of study; n = 184), the dominant bacterial taxa (Figure 1), as well as the alpha (Figure 2) and beta (Figure 3) diversities of CON animals, remained relatively stable over each year’s four-month study period, showing consistency across the breeding season. Adding the pretreatment phase of the other two groups (baseline samples; n = 43) to the control group (see ‘untreated average’ in Figure 1), the bacterial gut microbiota of healthy male ring-tailed lemurs, in captivity, were dominated by taxa in the Bacteroidetes and Firmicutes phyla, with lesser contributions from Proteobacteria, Spirochaetes, and Tenericutes. Within these five phyla, 20 genera accounted for minimally 1% of the total sequences (Figure 1).

**Figure 1.**
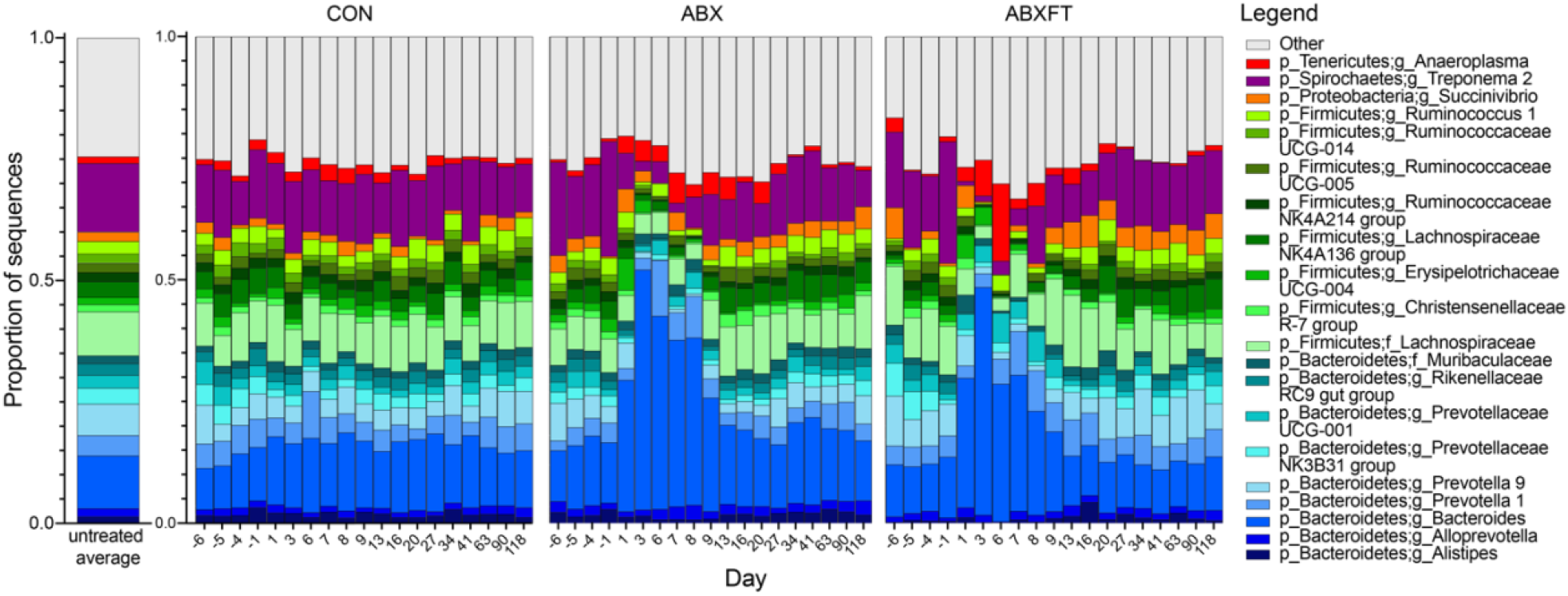
Mean relative abundances of bacterial genera over time in the gut microbiomes of three experimental groups of male ring-tailed lemurs (*Lemur catta*). Shown are values for healthy animals that received no treatment (CON), antibiotics only (ABX), or antibiotics plus fecal transfaunation (ABXFT). Genera are identified by color; those representing < 1% of the microbiomes were combined into the category “Other”. The x axis shows day relative to three phases of study: pretreatment (days −6 to −1), treatment (day 0-6/7), and recovery (day7/8-118).

**Figure 2.**
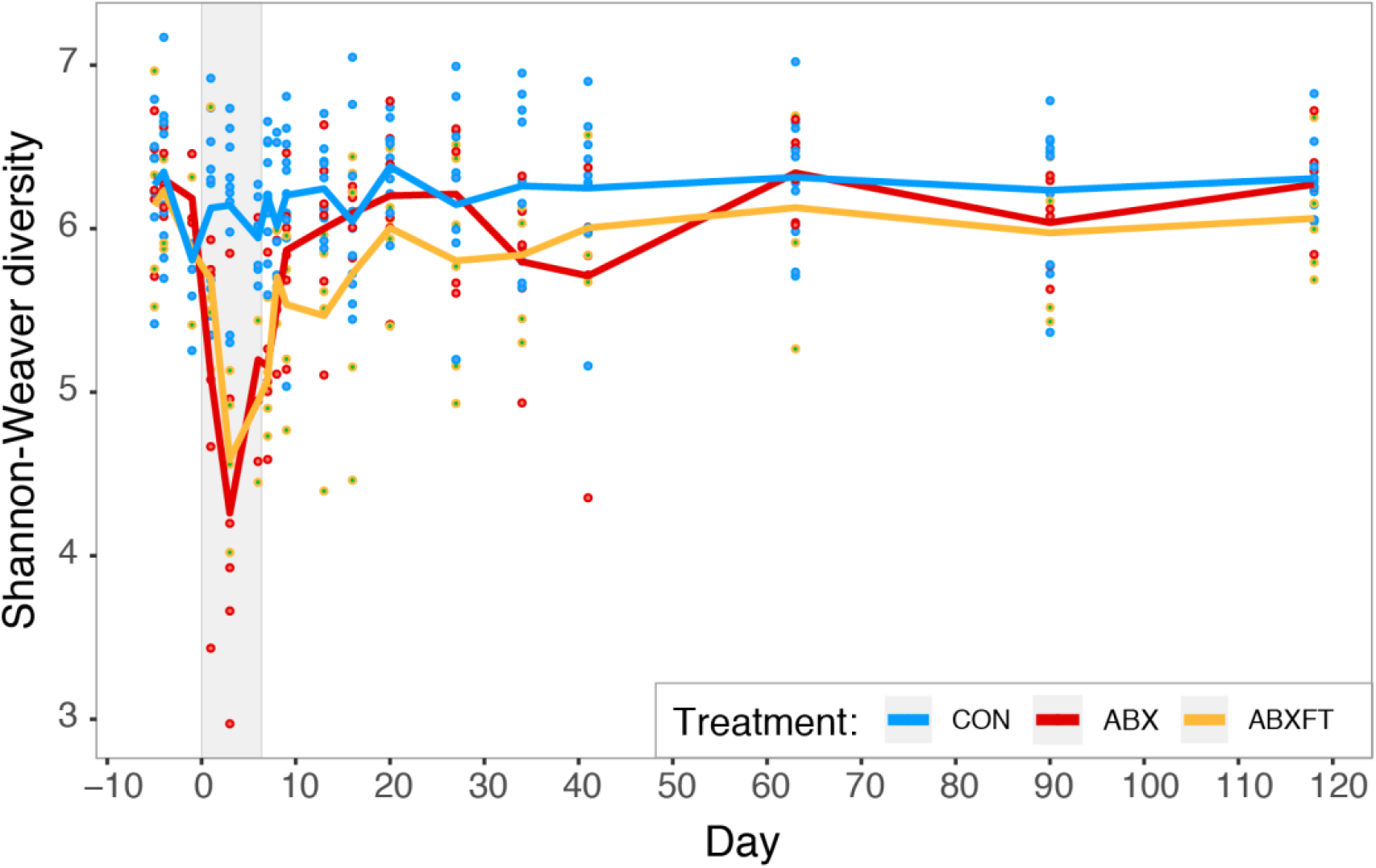
Shannon-Weaver alpha diversity over time in three experimental groups of male ring-tailed lemurs (*Lemur catta*). Shown are values for healthy animals that received no treatment (CON), antibiotics only (ABX), or antibiotics plus fecal transfaunation (ABXFT). Dots represent individual data points and lines connect the mean values of alpha diversity across individuals at each time point. The shaded window represents the period of antibiotic treatment (day 0-6), with fecal transfaunation administered on day 7; all values prior to the onset of treatment represent baseline values and all values post-treatment represent the period of recovery.

**Figure 3.**
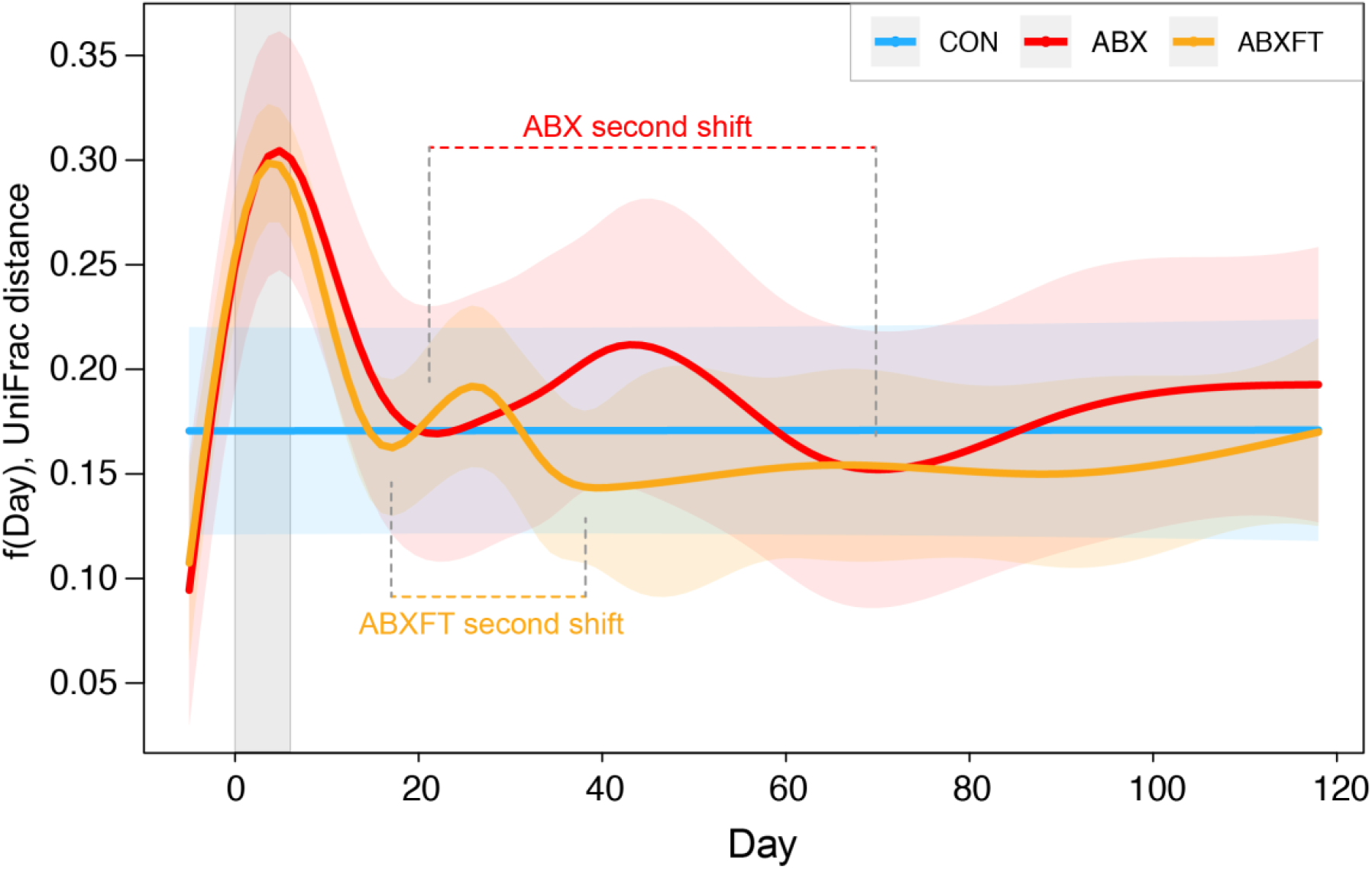
Beta diversity (Unweighted UniFrac distances) for three experimental groups of male ring-tailed lemurs (*Lemur catta*), represented as model-predicted distances from baseline, with 95% confidence intervals. Shown are values for healthy animals that received no treatment (CON), antibiotics only (ABX), or antibiotics plus fecal transfaunation (ABXFT). The gray shaded window represents the period of antibiotic treatment, with the prior period representing baseline and the subsequent period representing recovery. The second shifts away from baseline are identified and labelled for ABX and ABXFT animals.

### Response to and recovery from antibiotic treatment: alpha diversity and microbial membership

Across all three phases (day −6 – 120), we found a significant overall effect of experimental group on alpha diversity: relative to CON animals, ABX and ABXFT animals had significantly lower scores (HGAM: CON vs. ABX, t = −3.535, p < 0.001; CON vs. ABXFT, t = −4.007, p < 0.001; Figure 2). Neither year nor the interaction between year and experimental condition significantly related to bacterial alpha diversity (HGAM1: year, F = 0.001, p = 0.990; year*experimental condition, F = 0.942, p = 0.391), suggesting consistency of effects across both years.

As expected for the treatment phase, antibiotic-treated (ABX and ABXFT) animals showed a dramatic reduction in alpha diversity relative to CON animals (day 0-6; HGAM: CON vs. ABX, t = −4.534, p < 0.001; CON vs. ABXFT, t = −3.754, p < 0.001; see shaded bar in Figure 2). Consistent with the broad effects of amoxicillin, antibiotic treatment in healthy lemurs was associated with dramatically reduced representation across a wide range of taxa, including numerous taxa in the Firmicutes phylum, such as members of the Clostridiales class (e.g. Ruminococcaceae and Lachnospiraceae families). Certain taxa, however, were markedly unaffected by antibiotic treatment, including the *Bacteroides* genus and other members of the Bacteroidales family.

When focusing on the recovery phase only, we found that the differences in alpha diversity between CON and antibiotic-treated animals persisted over the nearly four-month, post-treatment period, suggesting long-lasting dysbiosis. Compared to CON animals, antibiotic-treated groups maintained significantly lower alpha diversity (HGAM: CON vs. ABX, t = −2.256, p = 0.025; CON vs. ABXFT, t = −3.036, p = 0.002); however, there was no significant differences between the alpha diversities of ABX and ABXFT animals during recovery (HGAM: ABX vs. ABXFT, t = 0.931, p = 0.354; Figure 2), suggesting no benefits of fecal transfaunation on alpha diversity. Unexpectedly, however, we observed an initial, rapid increase in alpha diversity in both experimental groups, consistent with the ‘diversity begets stability’ hypothesis (Figure 2).

### Response to and recovery from antibiotic treatment: beta diversity

Across all three phases, we also found experimental condition to be a significant predictor of beta diversity (HGAM2: F = 5.625, p = 0.004; Figure 3), but in a manner that differed from the findings on alpha diversity. Notably, compared to CON animals, ABX, but not ABXFT animals, showed significantly greater distances from their baseline communities (HGAM: CON vs. ABX, t = 3.434, p < 0.001; CON vs. ABXFT, t = 1.726, p = 0.085). Nevertheless, during the treatment phase, specifically, and compared to CON animals, both groups of antibiotic-treated animals showed significantly greater distances from their baseline communities (HGAM: CON vs. ABX, t = - 3.847, p < 0.001; CON vs. ABXFT, t = - 3.761, p < 0.001). Therefore, the recovery trajectories of the two treated groups diverged post treatment.

Indeed, during the recovery phase, CON animals were significantly less distant from baseline when compared to ABX animals, but not when compared to ABXFT animals (HGAM: CON vs. ABX, t = 2.790, p = 0.005; CON vs. ABXFT, t = 0.599, p = 0.549), consistent with the ‘key-stone species’ hypothesis. Furthermore, when comparing the recoveries of the two antibiotic-treated groups, ABX animals had significantly greater distance from baseline compared to ABXFT animals (HGAM: t = 2.115, p = 0.036; Figure 3). The bacterial composition of ABX animals continued to oscillate throughout the course of the experiment, whereas in ABXFT animals, bacterial composition became relatively stable approximately 2 weeks after the treatment phase (Figure 3), further supporting the results reported above. Specifically, after the first compositional shift during the treatment period, the bacterial composition of both ABX and ABXFT animals underwent a second shift away from baseline during the recovery period; however, the magnitude and span of these secondary shifts differed between the ABX and ABFT groups (Figure 3).

### Bacterial associations

To characterize the bacterial covariations that underlie microbial dynamics in the lemurs’ gut, we used pairwise covariation analyses; we detected several strong covariations (ρ > 0.5 or ρ < −0.5; hereafter ‘associations’) between pairs of microbes within the lemurs’ gut microbiomes (Table S1). We investigated these associations under all three experimental conditions in two stages: across all experimental phases and during the recovery phase.

Minimal variation within the microbiota of CON animals limited the detectability of normal bacterial associations; nevertheless, two strong associations emerged. The first was between the genus *Cerasicoccus* and the order WCHB1-41 and was evident across all experimental phases; the second was between the genus *Cerasicoccus* and the order Rhodospirillales, and was evident during the recovery phase (Table S1). These two relationships reflect the small-scale, yet ever-present, microbial dynamics that occur in healthy, unperturbed microbiomes.

Within the more variable gut microbiota of ABX and ABXFT lemurs (Figure 3), and across all three experimental phases, there were 35 and 31 strong associations, respectively (Figures 4a,b). In ABX animals, these associations were predominately positive, with only six negative associations, whereas in ABXFT lemurs, positive and negative associations were equally represented (15:16, respectively). Shared across ABX and ABXFT animals were 10 strong associations, eight positive and two negative. Within these shared associations, nine involved either *Parabacteroides* or *Bacteroides* (genus 12 and 39, respectively, in Figures 4a,b). The positive association between these two taxa was the strongest association for both ABX and ABXFT animals (Table S1). Moreover, in ABX and ABXFT animals, the log ratios of *Parabacteroides* and *Bacteroides* abundances both showed increases during the treatment phase, indicating that both taxa were relatively unaffected by antibiotic treatment (Figure 5). The majority of strong pairwise associations with *Parabacteroides* or *Bacteroides* were positive, indicating that the associated taxa also withstood the effects of antibiotic treatment, potentially via shared ABR genes.

**Figure 4.**
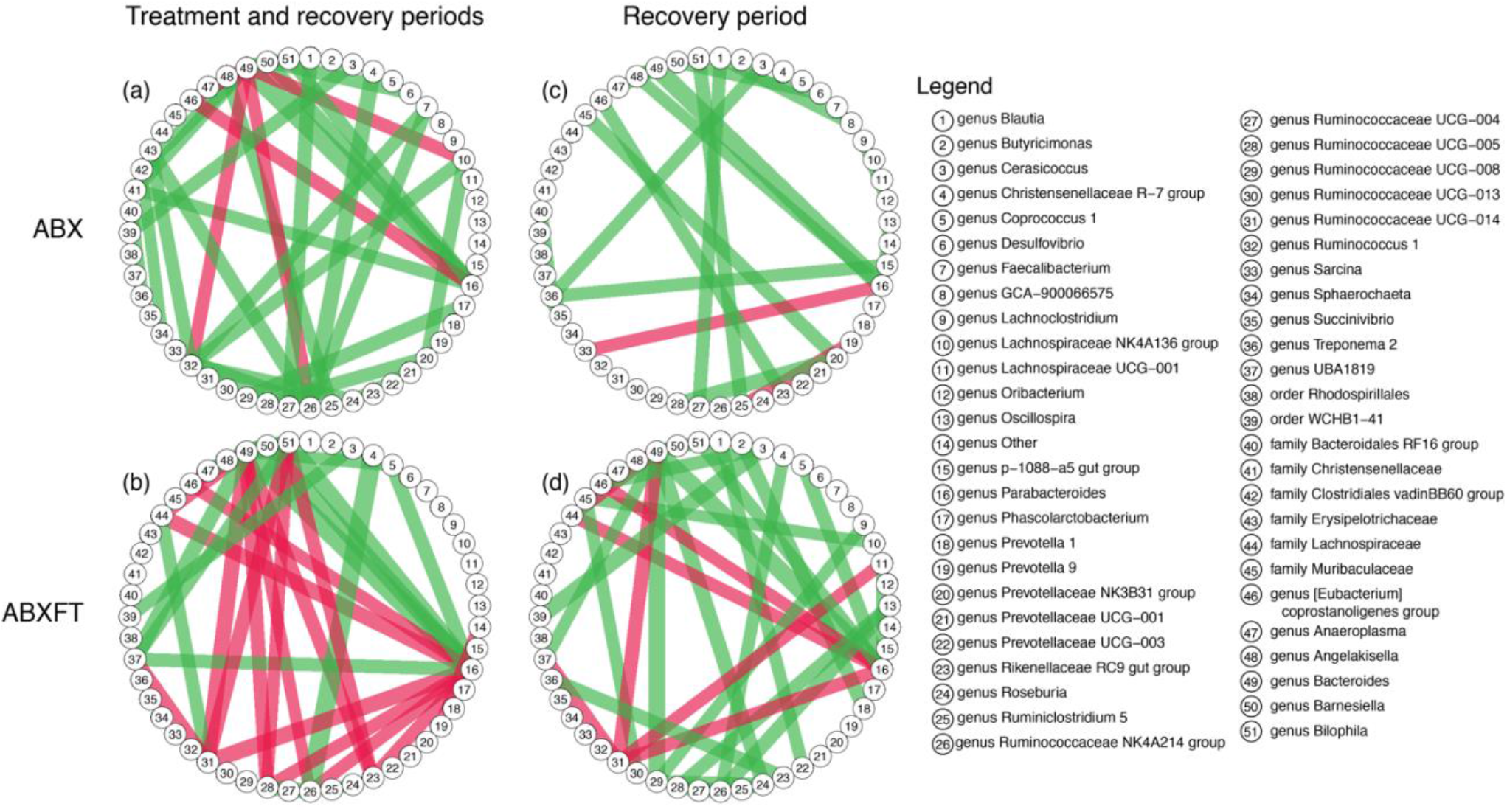
Bacterial associations for healthy, male ring-tailed lemurs (*Lemur catta*) either treated with antibiotics only (ABX) or with antibiotics plus a fecal transfaunation (ABXFT). Line colors represent the direction of the correlation (green = positive, red = negative); line width is scaled to the magnitude of the correlation.

**Figure 5.**
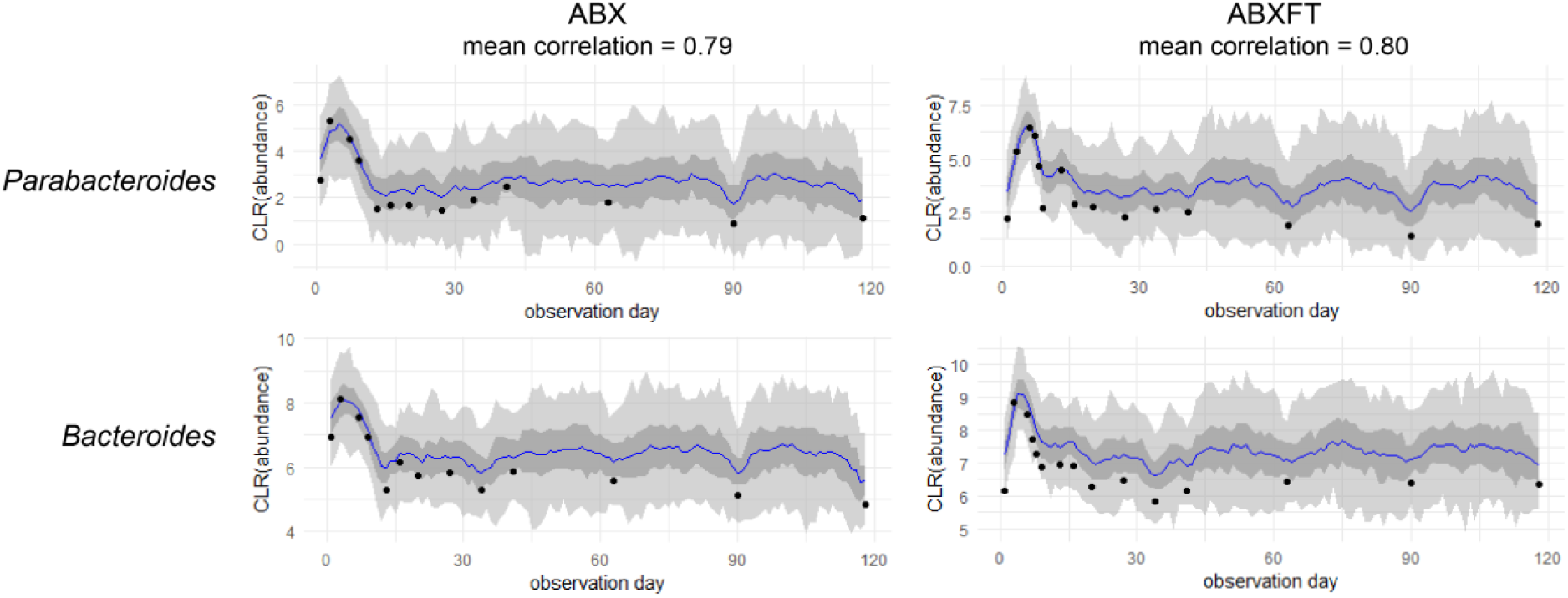
Representative correlation plots for the association between the Centered Log Ratio (CLR) of Bacteroides and Parabacteroides abundances in healthy, male ring-tailed lemurs (*Lemur catta*) either treated with antibiotics only (ABX) or with antibiotics plus fecal transfaunation (ABXFT). The antibiotic treatment period spans day 0-6, fecal transfaunation was administered on day 7, and all days thereafter constitute the period of recovery.

During the recovery phase, the gut microbiomes of ABX animals retained only 20 (18 positive and 2 negative) of the original 35 strong associations (Table S2, Figure 4c), likely reflecting the paucity of microbes that survived antibiotic treatment. By contrast, the gut microbiomes of ABXFT animals retained the same 31 (23 positive and 8 negative) strong associations (Table S2; Figure 4d), likely reflecting the reintroduction of baseline microbes, and their associations, through fecal transfaunation. Only two associations were shared between ABX and ABXFT animals during recovery: *Parabacteroides* and *Bacteroides*, and *Christensenellaceae R-7 group* and *Ruminococcaceae NK4A214 group*, both of which were also shared during the entire experimental period. Despite variability across treatment groups and phases, some of the strongest associations persisted during recovery (Table S2), further supporting the ‘key-stone species’ hypothesis.

### Cross-sectional and longitudinal ABR

Across the 30 fecal samples selected for shotgun sequencing (from a subset of subjects), 3.2 million sequences were assigned to 83 known ABR genes. On average, the majority of the ABR genes detected belonged to four resistance gene families: Tetracycline (51.4%), Beta-lactam (29.5%), Aminoglycoside (7.9%), and Macrolide (1.2%). There was also minor (>1%) representation of genes in the Vancomycin, Multi-Drug Resistant, and Sulphonamide families.

The cross-sectional data on the six, focal animals revealed unexpected variation in ABR. Notably, the two animals (IDs 7143 and 7086) that had never been treated with antibiotics nevertheless harbored ABR at levels similar to those of animals that had previously received numerous courses of antibiotics (Figure 6). Additionally, the animal (ID 6440) with the most numerous antibiotic treatments (n = 27 courses), harbored the second lowest abundance of ABR genes, similar to that of the animals with no previous treatment (Figure 6).

**Figure 6.**
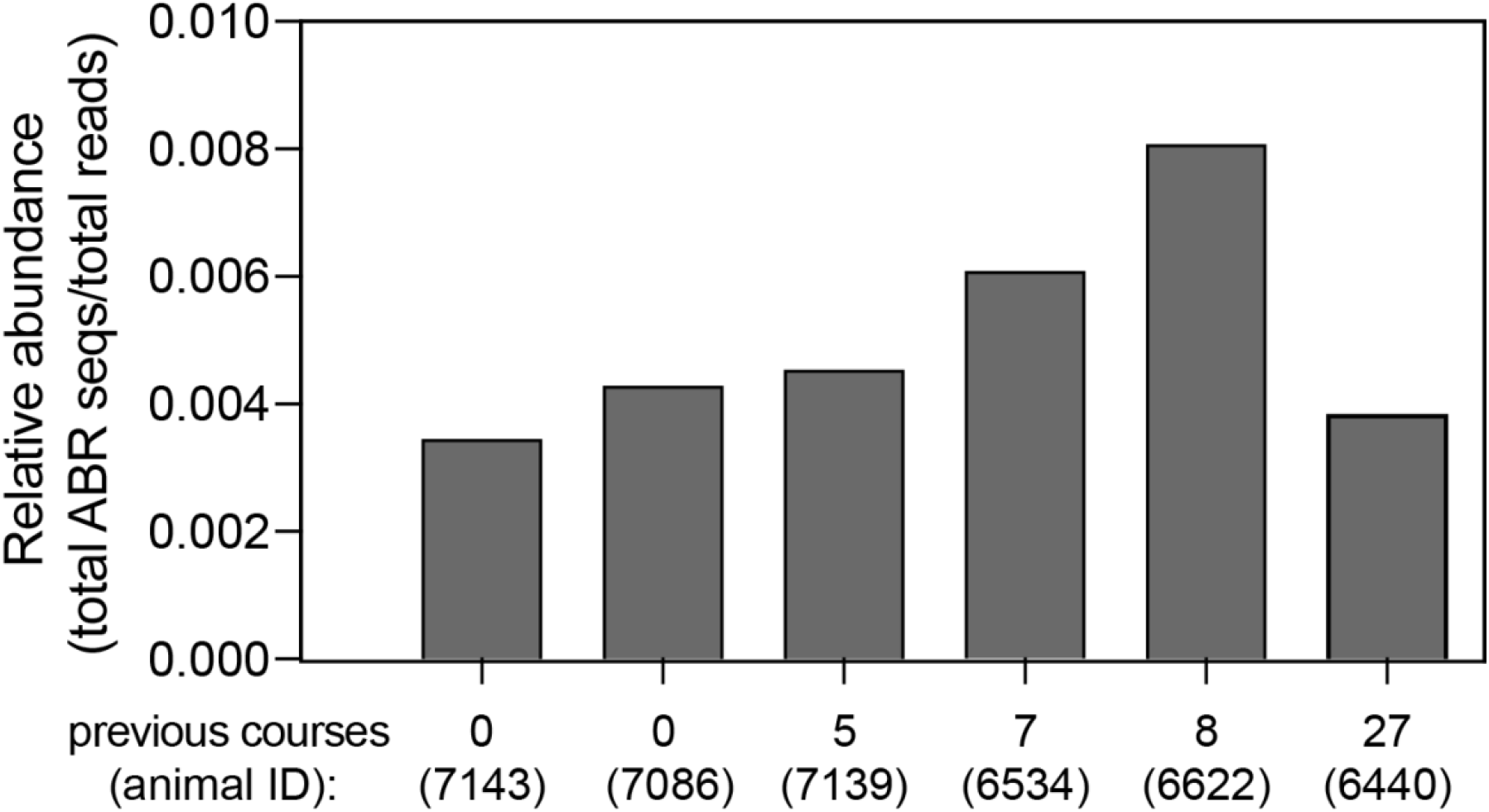
Relative abundance of antibiotic resistance (ABR) genes in six healthy, male ring-tailed lemurs (*Lemur catta*) that had received different numbers of treatment courses of antibiotics across their lifetime.

**Figure 7.**
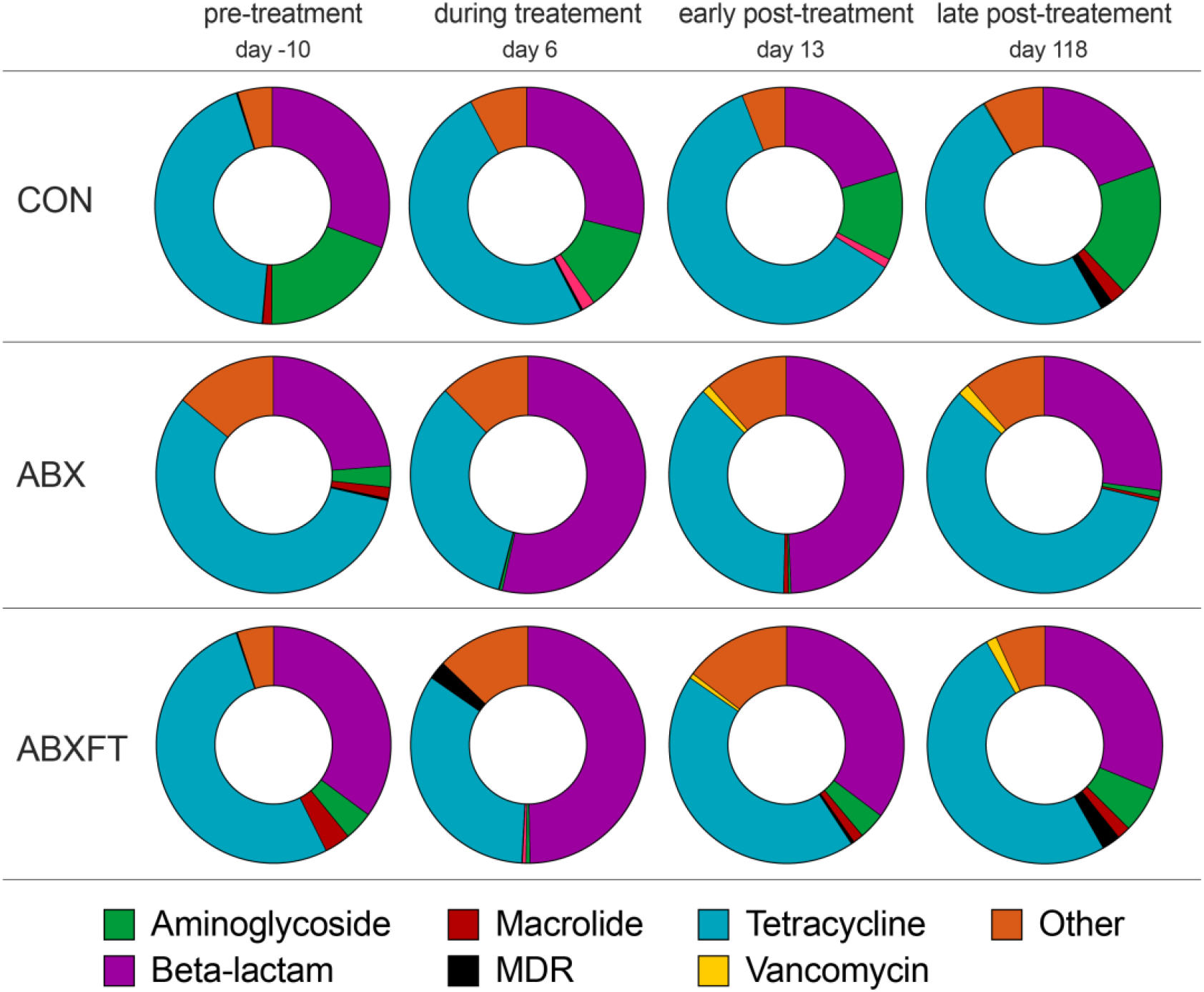
Proportions of antibiotic resistance (ABR) genes identified in healthy, male ring-tailed lemurs (*Lemur catta*) that received no treatment (CON), antibiotics only (ABX), or antibiotics plus fecal transfaunation (ABXFT). Shown are color-coded resistance gene families at four time points during the study, during which antibiotic treatment was administered on days 0-6 and fecal transfaunation was administered on day 7. MDR = Multi-Drug Resistant.

In analyses of longitudinal variation in the relative abundance of ABR genes, we found no significant effects of experimental condition or time point (HGAM: experimental condition, F = 0.530, p = 0.603; time point, F = 0.602, p = 0.627). By contrast, experimental condition significantly affected the proportion of ABR genes assigned to the beta-lactam resistance family. Compared to CON animals, ABX and ABXFT animals harbored significantly greater proportions of beta-lactam resistance genes (HGAM: CON vs. ABX, t = 2.352, p = 0.040, CON vs. ABXFT, t = 2.439, p = 0.034), reflecting the impact of treatment with a beta-lactam antibiotic (amoxicillin). An animal’s number of previous antibiotic courses was significantly, but negatively, related to the proportion of beta-lactam genes (HGAM: t = −2.766, p = 0.019). In ABX and ABXFT animals, samples collected pretreatment and late post-treatment had significantly fewer ABR genes than did samples collected during treatment (HGAM: during vs. pretreatment, t = −4.616, p = 0.003; during vs. late post-treatment, t = −4.605, p = 0.006). Although not statistically significant, the proportion of beta-lactam resistance genes during early post-treatment tended to be greater in ABX animals compared to ABXFT animals (HGAM: period and experiment group interaction effect, t = 1.766, p = 0.127; Figure 8), which may hint at a potential mitigating effect of fecal transfaunation on the persistence of ABR in lemur gut microbiomes.

**Figure 8.**
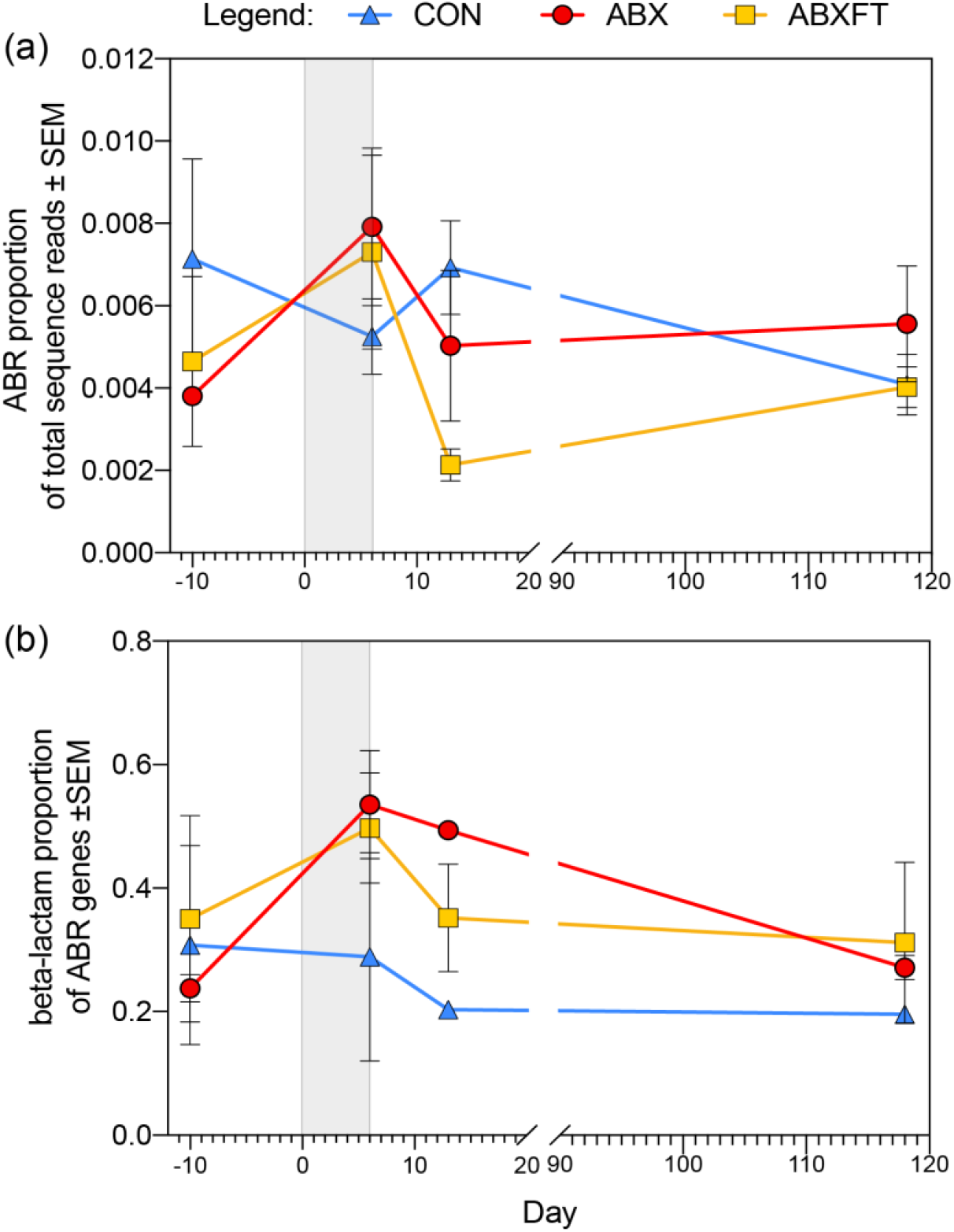
Patterns in antibiotic resistance across healthy, male ring-tailed lemurs (*Lemur catta*) that received no treatment (CON), antibiotics only (ABX), or antibiotics plus fecal transfaunation (ABXFT). Shown is variation over time in (a) the relative abundance of antibiotic resistance (ABR) genes and (b) the proportion of ABR genes assigned to the beta-lactam resistance gene family. The gray shaded window represents the period of antibiotic treatment, with the prior period representing baseline and the subsequent period representing recovery. SEM = standard error of the means.

## Discussion

In our longitudinal experiment, we contribute to the ecological framework for interpreting the dynamics of host-associated microbiomes by tracking bacterial recovery following antibiotic-mediated dysbiosis in healthy nonhuman primates. We provide support for both the ‘diversity begets stability’ and ‘key-stone species’ hypotheses, but with different schedules: microbial alpha diversity rebounded quickly (albeit incompletely) in treated animals, whereas beta diversity reflected a trajectory of long-term, microbial instability in animals that received antibiotics alone. The effects of fecal transfaunation on recovery of alpha diversity were negligeable, but for beta diversity, this procedure hastened and stabilized the recovery of community composition, supporting fecal transfaunation as an effective tool for treating microbial dysbiosis. The associations between bacteria varied between experimental conditions, suggesting that the relationships between microbial groups may have contributed to the differential effects of ABX and ABXFT treatments, which could have been governed by the presence of ABR genes. Our cross-sectional analysis of ABR showed that the prevalence of ABR genes in a host is not necessarily correlated with that host’s previous exposure to antibiotics; ABR can be acquired and maintained in the gut microbiomes of lemurs that had no previous antibiotic treatment. Longitudinally, ABR gene profiles reflected the type of antibiotic being used. As expected, the proportion of ABR genes that confer resistance to beta-lactamase antibiotics increased during treatment with amoxicillin. Lastly, fecal transfaunation may mitigate the persistence of ABR during recovery from antibiotic treatment. Using a holistic and longitudinal approach, across scales, allowed elucidating microbial dynamics that otherwise would have been imperceptible.

Consistent with previous findings^56,57^, and with the known efficacy of amoxicillin as a broad-spectrum antibiotic^58^, we found that animals receiving antibiotics concurrently showed a drastic decrease in alpha diversity. Contrary to expectations, however, lemurs post treatment showed a rapid rebound in alpha diversity, with ABX and ABFT animals showing no significant difference in their recovery trajectories. The rapidity of these patterns may owe to the healthy status of the hosts and to the relatively short period of antibiotic treatment. The ability to recolonize or re-diversify a microbiome after dysbiosis can be severely dampened by injury or disease^59,60^ and by recurrent antibiotic treatment^61,62^. Or, as omnivores with a broad dietary range, ring-tailed lemurs may have shown more rapid recovery than would be observed by dietary specialists ^42^. Beyond external influences (e.g. from diet, the physical environment, or social interaction), alpha diversity can also increase from within. Indeed, even if antibiotic treatment causes local bacterial extirpations, certain taxa can persist either by expressing ABR genes^8,63^ or by sequestering in areas of the gastrointestinal tract that are less affected by antibiotics (i.e., the appendix or cecum^64,65^), allowing for in-kind recolonization after disruption. Nevertheless, as evidenced by the results on beta diversity, early recovery of alpha diversity did not entail strictly in-kind recolonization of lemur gut microbiota. The rapidity in these patterns could thus lend support to the ‘diversity begets stability’ hypothesis, in that the key first step in community restoration may be to regain diversity, regardless of microbial composition.

Patterns of beta diversity elucidated longer-term effects of antibiotics on gut microbiome community composition. Throughout the four-month recovery period of ABX animals, the microbial community did not return to the baseline composition or even reach an alternative stable state. In line with previous evidence and with the key-stone species hypothesis, this pattern indicated that recouping key microbial members may be more elusive (and perhaps more critical to stability) than recouping sheer numbers of taxa^55^. Indeed, as predicted under this hypothesis, fecal transfaunation showed a stabilizing effect on community composition. The bacterial associations that characterized the recovery phase in ABXFT animals were more numerous and almost wholly different from those in ABX lemurs, suggesting that multitudinous microbial interactions underpin some of the effects of fecal transfaunation. These findings are consistent with the concept of competitive exclusion, whereby the diverse group of reintroduced, native bacteria outcompete pathogenic or opportunistic microbes^66,67^. Although there is much to learn about the modes of action in successful transfaunation, we add to the mounting evidence that they are a promising tool to hasten recovery from dysbiosis^68–70^.

Of the bacterial associations present in the two treatment conditions, *Parabacteroides* and *Bacteroides* – two, closely related taxa with similar functional potential ^71^ – dominated the observed relationships. The strength and quantity of relationships between *Bacteroides* and other bacterial taxa support the foundational membership of *Bacteroides* in the lemur gut microbiome^42,72,73^. Notably, increases in the log ratio abundance of *Bacteroides* during antibiotic treatment indicated that its members maintained or increased their relative abundances while other taxa were eliminated. Indeed, the *Bacteroides* genus is notorious for showing ABR. The diversity of its resistance mechanisms^74,75^, coupled with extensive lateral gene transfer within members of the genus and with *non-Bacteroides* taxa^76,77^, contributed to certain *Bacteroides spp*. having one of the highest resistance rates among known anaerobic pathogens^78^. Furthermore, certain *Bacteroides* harbor an unknown molecular mechanism that confers resistance specifically to amoxicillin^79^. Certain *Bacteroides* strains can even shield other taxa from the effects of beta-lactam antibiotics^80^. Combined with this evidence, our results suggest that *Bacteroides* in the lemur gut microbiome likely have amoxicillin resistance mechanisms that enable them to dictate most bacterial relationships during recovery from antibiotic-induced dysbiosis.

ABR genes, including some that are considered clinically relevant^81^, were present within the gut microbiome of all lemurs. Somewhat surprisingly, lemurs that had never received antibiotic treatment showed resistance levels similar to those of lemurs previously treated with antibiotics. Researchers have shown that bacteria and their genes can be shared between hosts that cohabitate^82,83^ or share social partners^84^. Furthermore, ABR genes often reside on mobile genetic units and are prone to rapid transfer between microbes^85^. Indeed, the ‘resistance crisis’ is perpetuated by the ubiquitous spread of ABR genes around the world^10,86,87^. Here, we find that lemurs are not exempt from these phenomena and, for captive animals especially, ABR could pose a severe threat to animal health^88,89^. Methods to mitigate the development and spread of ABR among animal populations, including perhaps via fecal transfaunation, may prove to be a critical facet of combatting the resistance crisis^86,90^.

Collectively, these results further our understanding of host-microbe relationships in the Anthropocene era ^91,92^. Evolutionary medicine is founded on the principle that, over evolutionary and proximate scales, changing environments and conditions influence aspects of well-being^93–95^. As scientists and medical practitioners increasingly recognize the value of a One Health perspective on human medicine (i.e., that human, animal, and environmental health are inseparable), holistic approaches will add context to our understanding of clinically relevant phenomena (i.e., the effects of antibiotics and ABR; ^87,96,97^). Ultimately, shedding light on how ‘old friends’ react to aspects of the ‘new world’ is relevant both to our understanding of the evolution of symbiosis and to its implications for modern medicine.

## Methods

### Study subjects and housing

Our study subjects were 11 healthy, reproductively intact, adult (4-16 yrs old), male ring-tailed lemurs housed in 10 conspecific, mixed-sex groups at the Duke Lemur Center (DLC; Durham, NC, USA). Within a two-year period, 10 of them underwent a control round with no treatment, but all 11 underwent one round of antibiotic treatment (see below), while living in their same social groups. During inclement weather (<5 °C), the groups would be sequestered in indoor enclosures, otherwise, they all had access to indoor and outdoor enclosures (approximately 146 m^2^/animal). Some of the groups additionally had access to large, forest enclosures where they semi-free-ranged with heterospecific lemurs. The animals received a diet of produce and commercial primate chow and, while semi-free-ranging, had access to natural foods foraged from the forest. Additional information on the lemurs’ diet, foraging, and social behavior have been reported elsewhere ^98^. The subjects were maintained in accordance with the NIH Guide for the Care and Use of Laboratory Animals, and procedures were approved by the Institutional Animal Care and Use Committee of Duke University (protocol A111-16-05).

### Study design and sample collection

To allow for a partial cross-over design, we conducted the study during two matched periods (October-February) in consecutive years, during the subjects’ breeding season in the Northern Hemisphere ^99^: 2016-2017 (Y1, n = 10 subjects) and 2017-2018 (Y2, n = 11 subjects). In each year, we assigned the subjects to one of three experimental groups: control animals (CON; Y1, n = 4; Y2, n = 6), antibiotic-treated animals (ABX; Y1, n = 3; Y2, n = 3), and antibiotic-treated animals receiving a fecal transfaunation (ABXFT; Y1, n = 3; Y2, n = 2). To avoid administering antibiotics twice to any animal, each animal was assigned to the CON group in one of the two years.

Each year of study involved three phases, lasting a total of ~125 days: a pretreatment or baseline phase (lasting ~ 6 days; i.e., day −6 to −1), a treatment phase (lasting 7-8 days; i.e., day 0 to 6/7), and a recovery phase (lasting ~110 days). In the treatment phase, all treated animals (ABX and ABXFT; n = 11) received a 7-day course of the broad-spectrum, beta-lactam antibiotic, amoxicillin (10 mg/kg body weight, received orally, twice daily). Approximately 24 hrs after completion of the full antibiotic regimen, ABXFT subjects received a fecal transfaunation consisting of their own feces: 2-3 fecal pellets, collected pretreatment, were mixed with water and administered orally via syringe or feeding tube, according to routine procedures that have been adopted by the DLC since the mid 1980s to treat outbreaks of gastrointestinal diseases ^45,100,101^.

The phases were additionally differentiated by the frequency with which we collected fecal samples: We collected samples every 1-3 days before, during, and immediately after the treatment phase, after which sampling occurred every 5-28 days. Typically, upon the subject’s morning voiding, between 7:00 am and 11:30 am, we opportunistically collected fresh fecal samples. On occasion, we collected samples from awake, gently restrained animals that were habituated to capture and collection procedures. At each time point, we sampled all subjects and we maintained analogous sampling regimes across years. We collected all samples in sterile, 15-ml falcon tubes, immediately placed them on ice, and then stored them in a −80 °C freezer within 2 hours, until analysis.

### Microbial DNA extraction, sequencing and bioinformatics

Using the DNeasy Powersoil kit (QIAGAN, Frederick, MD), we extracted microbial gDNA from fecal samples and from four blank controls, to control for possible contamination. We quantified the extracted gDNA using a Fluorometer (broad-spectrum kit, Qubit 4, Thermo Fisher Scientific, Waltham, MA). These extractions were used for bacterial identification (via 16S rRNA amplicon sequencing) and ABR gene identification (via shotgun sequencing), as described below.

### Bacterial identification

We shipped aliquots of extracted gDNA to the Argonne National Laboratory’s Environmental Sequencing facility (Lemont, IL) for library preparation and sequencing of the 16S rRNA gene. There, the V4 region of the 16S rRNA gene (515F-806R) was amplified with region-specific primers adapted for the Illumina MiSeq platform ^102^. Forward primers contained a 12-base barcode sequence to support pooling of samples in each flow cell lane. Once pooled, amplicon libraries were cleaned using AMPure XP Beads (Beckman Coulter, Pasadena, CA), and quantified using a fluorometer (Qubit 4). Amplicons were sequenced on a 2 x 151 bp Illumina MiSeq run ^102^. Sequencing reads are available on the National Center for Biotechnology Information’s Sequence Read Archive (BioProject ID #TBD, BioSample accession #s TBD).

In collaboration with Duke University’s Genomic Analysis and Bioinformatics Shared Resource, 16S raw sequence data were analyzed using a bioinformatics pipeline generated in QIIME2 (ver 2018.11) ^103^. The pipeline included steps to join, demultiplex, and quality-filter sequence reads. The DADA2 plugin (q2-dada2) ^104^ was used to denoise, quality-filter, and remove phiX and chimeric sequences from the demultiplexed reads. Using the resulting sequences, we discarded samples with < 10,000 reads. To determine taxonomic classification, we used a pre-trained Naive Bayes classifier at 99% sequence identity (SILVA-132 database) ^105,106^. After bioinformatic processing, a total of 344 fecal samples (from all subjects across all study phases) yielded over 23.4 million 16S sequences (mean per sample = 59,766).

We used the feature tables and taxonomies of bacterial members to calculate Shannon-Weaver diversity (i.e., alpha diversity). To assess microbial composition (i.e., beta diversity), we calculated unweighted UniFrac, a distance metric that considers the phylogenetic relationships between taxa. After calculating these diversity metrics, we combined features without assigned taxonomy below the Kingdom level into an “Unassigned” category. We also included the conglomerate “Other” to represent the rare taxa that had relative abundances < 1%.

### ABR identification

To allow for cross-sectional and longitudinal analyses of ABR genes, while limiting the expense of metagenomic analyses, we performed shotgun sequencing on samples from six lemurs, two per experimental group, that ranged in their previous exposure to antibiotics (0-27 previous courses). For cross-sectional analysis, we included one sample from each subject’s pretreatment phase in Y1. For longitudinal analysis (which we prioritized), we included four samples from each animal (days −10, 6, 13, and 118) in Y2. We shipped this subset of extractions (n = 30) for shotgun sequencing to CosmosID (Rockville, MD), where DNA libraries were prepared using the Illumina Nextera XT library preparation kit, with a modified protocol. Library quantity was assessed with Qubit (ThermoFisher). Libraries were then sequenced on an Illumina HiSeq platform 2 x 150 bp.

The samples selected for shotgun sequencing averaged 20.4 million sequences per sample. The resulting unassembled sequencing reads underwent multi-kingdom microbiome analysis and profiling of antibiotic resistance genes using the CosmosID bioinformatics platform (CosmosID Inc., Rockville, MD), as described elsewhere ^107,108^. The antibiotic resistance and virulence genes in the microbiome were identified by querying the unassembled sequence reads against the CosmosID-curated antibiotic resistance and virulence associated gene databases ^109,110^.

### Statistical analyses

To characterize variation in bacterial diversity and composition, we used Hierarchical General Additive Models (HGAM) ^111^, which have the flexibility to accommodate nonlinear trends (See Supplementary Materials 1 for full model syntax). We used this model to test for patterns in bacterial alpha and beta diversity. For analyses of beta diversity, we first used Principal Coordinates Analysis (PCoA) to visualize variation in bacterial composition (UniFrac distance) in coordinate space. Subsequently, we used distance metrics to calculate change in bacterial composition relative to a pretreatment, baseline community (collected 4 days before the onset of treatment for all animals; QIIME2). We tested for variation in these calculated distance measures using our HGAM. To assess the response to and recovery from treatment, we first used our models to test for variation across the entire dataset and, then, using subsets of the alpha and beta diversity data, we focused our model on the treatment and recovery phases.

To better understand the short- and longer-term process of recovery, we tested for associations between bacterial taxa over time and evaluated how these associations may have differed between treatment groups. To exclude spurious associations, we first removed a few pretreatment samples from each animal’s series, further allowing us to focus on associations during the treatment and recovery phases. To reduce sparsity in the dataset and ease the computational burden, we removed rare taxa present in < five samples across the full dataset, clustered taxa at the genus level, and grouped as ‘Other’ all low-abundance genera with < 0.01% of total counts. This filtering removed < 1% of total sequence counts.

To naturally model the irregular temporal spacing in the observations and manage autocorrelation between samples, we fitted a Bayesian multivariate Gaussian process to each of multiple synthetic replicates of the dataset (see resampling procedure below). We then inferred a distribution over the covariance between microbes. The sample collection schedule motivated two key choices in noise modeling and data representation. First, because stochasticity exists in sample collection, processing, and sequencing, we used a resampling method similar to that of ALDEx2^112,113^ to emulate the variation that would be expected from replicate measurements. Second, to account for the compositional nature of the sequence count data within our model ^114^, we used log ratios as compositional representations of the data. We converted the estimated covariance matrices to correlation matrices and thresholded all pairwise correlations between microbes to select as significant those with 95% credible intervals that excluded zero correlation (i.e., those with strong positive or negative associations). We then ranked associations by their median strength and selected those in excess of correlation ρ > 0.5 or ρ < −0.5 as strong associations.

Because our cross-sectional ABR data had small sample sizes and minimal statistical power, we were limited to examining qualitative trends. To assess patterns in longitudinal ABR data, we used another HGAM model (Supplementary Materials 1), fitted to the following aspects of ABR gene profiles: Across all three experimental groups, we tested for patterns in the relative abundance of all sequences that were assigned to an ABR gene and in the proportion of ABR genes assigned to beta-lactam resistance genes. To further investigate the effects of antibiotics and fecal transfaunation on ABR, we ran additional HGAMs on the ABR data from only the ABX and ABXFT groups.

## Acknowledgements

We thank the current and past staff members of the DLC, particularly David Brewer, Erin Ehmke, Megan Davison, Cat Ostrowski, Bobby Schopler, Melanie Simmons, and Kay Welser, for their insights and help with animal handling. Conversations with Lydia Greene were instrumental during this project’s inception. We are grateful to Sarah Owens at Argonne National Laboratory and Karlis Graubics and Brian Fanelli at CosmosID provided guidance and sequencing services. David Corcoran and Zhengzheng Wei at the Duke Genomic Analysis and Bioinformatics Shared Resource provided bioinformatics analyses. Funding was provided by awards from the National Science Foundation (BCS 1749465 to CD), DLC Director’s Fund (to SB and RH), and Duke Microbiome Center (formerly Duke Center for the Genomics of Microbial Systems, Grants-In-Aid for Microbiome Bioinformatic Analysis to CD). This is DLC publication number (##TBD).

## References

1 Koskella B, Bergelson J. The study of host–microbiome (co) evolution across levels of selection. Philos Trans R Soc B 2020; 375: 20190604.

2 Lynch JB, Hsiao EY. Microbiomes as sources of emergent host phenotypes. Science (80-) 2019; 365: 1405–1409.

3 Brown K, DeCoffe D, Molcan E, Gibson DL. Diet-induced dysbiosis of the intestinal microbiota and the effects on immunity and disease. Nutrients 2012; 4: 1095–1119.

4 Li J, Zhao F, Wang Y, Chen J, Tao J, Tian G, Wu S, Liu W, Cui Q, Geng B. Gut microbiota dysbiosis contributes to the development of hypertension. Microbiome 2017; 5: 1–19.

5 Moloney RD, Desbonnet L, Clarke G, Dinan TG, Cryan JF. The microbiome: stress, health and disease. Mamm Genome 2014; 25: 49–74.

6 Rook GAW. 99th Dahlem conference on infection, inflammation and chronic inflammatory disorders: Darwinian medicine and the ‘hygiene’or ‘old friends’ hypothesis. Clin Exp Immunol 2010; 160: 70–79.

7 Rook GAW, Martinelli R, Brunet LR. Innate immune responses to mycobacteria and the downregulation of atopic responses. Curr Opin Allergy Clin Immunol 2003; 3: 337–342.

8 Francino MP. Antibiotics and the human gut microbiome: dysbioses and accumulation of resistances. Front Microbiol 2016; 6: 1–11.

9 Langdon A, Crook N, Dantas G. The effects of antibiotics on the microbiome throughout development and alternative approaches for therapeutic modulation. Genome Med 2016; 8: 39.

10 Ventola CL. The antibiotic resistance crisis: part 1: causes and threats. Pharm Ther 2015; 40: 277.

11 Taur Y, Coyte K, Schluter J, Robilotti E, Figueroa C, Gjonbalaj M, Littmann ER, Ling L, Miller L, Gyaltshen Y et al. Reconstitution of the gut microbiota of antibiotic-treated patients by autologous fecal microbiota transplant. Sci Transl Med 2018; 10.

12 D’Costa VM, King CE, Kalan L, Morar M, Sung WWL, Schwarz C, Froese D, Zazula G, Calmels F, Debruyne R et al. Antibiotic resistance is ancient. Nature 2011; 477: 457–461.

13 Davies JE. Origins, acquisition and dissemination of antibiotic resistance determinants. In: Chadwick DJ, Goode JA (eds). Antibiotic Resistance: Origins, Evolution, Selection and Spread. Wiley Online Library, 1997, pp 15–27.

14 Atwood KC, Schneider LK, Rryan FJ. Selective mechanisms in bacteria. In: Cold Spring Harbor Symposia on Quantitative Biology. Cold Spring Harbor Laboratory Press, 1951, pp 345–355.

15 Denamur E, Matic I. Evolution of mutation rates in bacteria. Mol Microbiol 2006; 60: 8208–27.

16 Ochman H, Lawrence JG, Groisman E. Lateral gene transfer and the nature of bacterial innovation. Nature 2000; 405: 299–304.

17 Thomas CM, Nielsen KM. Mechanisms of, and barriers to, horizontal gene transfer between bacteria. Nat Rev Microbiol 2005; 3: 711–721.

18 Allen HK, Donato J, Wang HH, Cloud-Hansen KA, Davies J, Handelsman J. Call of the wild: antibiotic resistance genes in natural environments. Nat Rev Microbiol 2010; 8: 251–259.

19 Aminov RI. The role of antibiotics and antibiotic resistance in nature. Environ Microbiol 2009; 11: 2970–2988.

20 Ferrer M, Méndez-García C, Rojo D, Barbas C, Moya A. Antibiotic use and microbiome function. Biochem Pharmacol 2017; 134: 114–126.

21 Cho I, Yamanishi S, Cox L, Methe BA, Zavadil J, Li K, Gao Z, Mahana D, Raju K, Teitler I et al. Antibiotics in early life alter the murine colonic microbiome and adiposity. Nature 2012; 488: 621–626.

22 Zarrinpar A, Chaix A, Xu ZZ, Chang MW, Marotz CA, Saghatelian A, Knight R, Panda S et al. Antibiotic-induced microbiome depletion alters metabolic homeostasis by affecting gut signaling and colonic metabolism. Nat Commun 2018; 9: 1–13.

23 Kolář M, Urbánek K, Látal T. Antibiotic selective pressure and development of bacterial resistance. Int J Antimicrob Agents 2001; 17: 357–363.

24 Witte W. Selective pressure by antibiotic use in livestock. Int J Antimicrob Agents 2000; 16: 19–24.

25 Stokes HW, Gillings MR. Gene flow, mobile genetic elements and the recruitment of antibiotic resistance genes into Gram-negative pathogens. FEMS Microbiol Rev 2011; 35: 790–819.

26 Weber DJ, Raasch R, Rutala WA. Nosocomial infections in the ICU: the growing importance of antibiotic-resistant pathogens. Chest 1999; 115: 34S–41S.

27 Britton RA, Young VB. Interaction between the intestinal microbiota and host in *Clostridium difficile* colonization resistance. Trends Microbiol 2012; 20: 313–319.

28 Stecher B, Maier L, Hardt W-D. ’Blooming’in the gut: how dysbiosis might contribute to pathogen evolution. Nat Rev Microbiol 2013; 11: 277.

29 Chambers HF, DeLeo FR. Waves of resistance: *Staphylococcus aureus* in the antibiotic era. Nat Rev Microbiol 2009; 7: 629–641.

30 Petty NK, Zakour NL Ben, Stanton-Cook M, Skippington E, Totsika M, Forde BM, Phan M-D, Moriel DG, Peters KM, Davies M et al. Global dissemination of a multidrug resistant *Escherichia coli* clone. Proc Natl Acad Sci 2014; 111: 5694–5699.

31 Aroniadis OC, Brandt LJ. Fecal microbiota transplantation: past, present and future. Curr Opin Gastroenterol 2013; 29: 79–84.

32 Grehan MJ, Borody TJ, Leis SM, Campbell J, Mitchell H, Wettstein A. Durable alteration of the colonic microbiota by the administration of donor fecal flora. J Clin Gastroenterol 2010; 44: 551–561.

33 Bo T-B, Zhang X-Y, Kohl KD, Wen J, Tian S-J, Wang D-H. Coprophagy prevention alters microbiome, metabolism, neurochemistry, and cognitive behavior in a small mammal. ISME J 2020; 14: 2625–2645.

34 Osawa R, Blanshard WH, Ocallaghan PG. Microbiological studies of the intestinal microflora of the koala, *Phascolarctos cinereus*. 2. Pap, a special maternal feces consumed by juvenile koalas. Aust J Zool 1993; 41: 611–620.

35 Chaitman J, Jergens AE, Gaschen F, Garcia-Mazcorro JF, Marks SL, Marroquin-Cardona AG, Richter K, Rossi G, Suchodolski JS, Weese JS et al. Commentary on key aspects of fecal microbiota transplantation in small animal practice. Vet Med Res Reports 2016; 7: 71.

36 Niederwerder MC. Fecal microbiota transplantation as a tool to treat and reduce susceptibility to disease in animals. Vet Immunol Immunopathol 2018; 206: 65–72.

37 Bakken JS, Borody T, Brandt LJ, Brill J V, Demarco DC, Franzos MA, Kelly C, Khoruts A, Louie T, Martinelli LP et al. Treating *Clostridium difficile* infection with fecal microbiota transplantation. Clin Gastroenterol Hepatol 2011; 9: 1044–1049.

38 Brandt LJ, Aroniadis OC, Mellow M, Kanatzar A, Kelly C, Park T, Stollman N, Rohlke F, Surawicz C et al. Long-Term Follow-Up of Colonoscopic Fecal Microbiota Transplant for Recurrent *Clostridium difficile* Infection. Am J Gastroenterol 2012; 107: 1079–1087.

39 Amato KR, Sanders JG, Song SJ, Nute M, Metcalf JL, Thompson LR, Morton JT, Amir A, McKenzie VJ, Humphrey G et al. Evolutionary trends in host physiology outweigh dietary niche in structuring primate gut microbiomes. ISME J 2019; : 1.

40 Clayton JB, Gomez A, Amato K, Knights D, Travis DA, Blekhman R, Knight R, Leigh S, Stumpf R, Wolf T et al. The gut microbiome of nonhuman primates: Lessons in ecology and evolution. Am J Primatol 2018; : e22867.

41 Greene LK, Clayton JB, Rothman RS, Semel BP, Semel MA, Gillespie TR, Wright PC, Drea CM. Local habitat, not phylogenetic relatedness, predicts gut microbiota better within folivorous than frugivorous lemur lineages. Biol Lett 2019; 15: 20190028.

42 Greene LK, Bornbusch SL, McKenney EA, Harris RL, Gorvetzian SR, Yoder AD, Drea CM. The importance of scale in comparative microbiome research: New insights from the gut and glands of captive and wild lemurs. Am J Primatol 2019.

43 Gould L. Lemur catta Ecology : What We Know and What We Need to Know. In: Gould L, Sauther ML (eds). Lemurs: Ecology and Adaptation. Springer: New York, 2006, pp 255–274.

44 Jolly A, Sussman RW, Koyama N, Rasamimanana H. Ringtailed lemur biology: Lemur catta in Madagascar. Springer Science & Business Media, 2006.

45 McKenney EA, Greene LK, Drea CM, Yoder AD. Down for the count: *Cryptosporidium* infection depletes the gut microbiome in Coquerel’s sifakas. Microb Ecol Health Dis 2017; 28:1335165.

46 Konopka A. What is microbial community ecology? ISME J 2009; 3: 1223–1230.

47 Moya A, Ferrer M. Functional redundancy-induced stability of gut microbiota subjected to disturbance. Trends Microbiol 2016; 24: 402–413.

48 Wohl DL, Arora S, Gladstone JR. Functional redundancy supports biodiversity and ecosystem function in a closed and constant environment. Ecology 2004; 85: 1534–1540.

49 Johnson KH, Vogt KA, Clark HJ, Schmitz OJ, Vogt DJ. Biodiversity and the productivity and stability of ecosystems. Trends Ecol Evol 1996; 11: 372–377.

50 McNaughton SJ. Diversity and stability of ecological communities: a comment on the role of empiricism in ecology. Am Nat 1977; 111: 515–525.

51 Mccann KS. The diversity–stability debate. 2000; 405.

52 Banerjee S, Schlaeppi K, van der Heijden MGA. Keystone taxa as drivers of microbiome structure and functioning. Nat Rev Microbiol 2018; 16: 567–576.

53 Berry D, Widder S. Deciphering microbial interactions and detecting keystone species with co-occurrence networks. Front Microbiol 2014; 5: 219.

54 Fisher CK, Mehta P. Identifying keystone species in the human gut microbiome from metagenomic timeseries using sparse linear regression. PLoS One 2014; 9: e102451.

55 Gibbons SM. Keystone taxa indispensable for microbiome recovery. Nat Microbiol 2020; 5: 1067–1068.

56 Palleja A, Mikkelsen KH, Forslund SK, Kashani A, Allin KH, Nielsen T, Hansen TH, Liang S, Feng Q, Zhang C et al. Recovery of gut microbiota of healthy adults following antibiotic exposure. Nat Microbiol 2018; 3: 1255–1265.

57 Vlčková K, Gomez A, Petrželková KJ, Whittier CA, Todd AF, Yeoman CJ, Nelson KE, Wilson BA, Stumpf RM, Modrý D et al. Effect of antibiotic treatment on the gastrointestinal microbiome of free-ranging western lowland gorillas (*Gorilla g. gorilla*). Microb Ecol 2016; 72: 943–954.

58 Kaur SP, Rao R, Nanda S. Amoxicillin: a broad spectrum antibiotic. Int J Pharm Pharm Sci 2011; 3: 30–37.

59 Krezalek MA, Alverdy JC. The role of the microbiota in surgical recovery. Curr Opin Clin Nutr Metab Care 2016; 19: 347–352.

60 Nicholson SE, Merrill D, Zhu C, Burmeister DM, Zou Y, Lai Z, Darlington DN, Lewis AM, Newton L, Scroggins S et al. Polytrauma independent of therapeutic intervention alters the gastrointestinal microbiome. Am J Surg 2018; 216: 699–705.

61 McDonald LC. Effects of short-and long-course antibiotics on the lower intestinal microbiome as they relate to traveller’s diarrhea. J Travel Med 2017; 24: S35–S38.

62 Korpela K, Salonen A, Virta LJ, Kekkonen RA, Forslund K, Bork P, De Vos WM. Intestinal microbiome is related to lifetime antibiotic use in Finnish pre-school children. Nat Commun 2016; 7: 10410.

63 Van Schaik W. The human gut resistome. Philos Trans R Soc B Biol Sci 2015; 370: 20140087.

64 Bollinger RR, Barbas AS, Bush EL, Lin SS, Parker W. Biofilms in the normal human large bowel: fact rather than fiction. Gut 2007; 56: 1481–1482.

65 Laurin M, Everett M Lou, Parker W. The cecal appendix: one more immune component with a function disturbed by post-industrial culture. Anat Rec Adv Integr Anat Evol Biol 2011; 294: 567–579.

66 Willing BP, Jansson JK. The gut microbiota: ecology and function. Lawrence Berkeley National Lab, Berkeley, CA (United States), 2010.

67 Zeng W, Shen J, Bo T, Peng L, Xu H, Nasser MI, Zhuang Q, Zhao M. Cutting edge: Probiotics and fecal microbiota transplantation in immunomodulation. J Immunol Res 2019; 2019.

68 Jin Song S, Woodhams DC, Martino C, Allaband C, Mu A, Javorschi-Miller-Montgomery S, Suchodolski JS, Knight R. Engineering the microbiome for animal health and conservation. Exp Biol Med 2019; 244: 494–504.

69 Schmidt EKA, Torres-Espin A, Raposo PJF, Madsen KL, Kigerl KA, Popovich PG, Fenrich KK, Fouad K. Fecal transplant prevents gut dysbiosis and anxiety-like behaviour after spinal cord injury in rats. PLoS One 2020; 15: e0226128.

70 Wei Y, Gong J, Zhu W, Guo D, Gu L, Li N, Li J. Fecal microbiota transplantation restores dysbiosis in patients with methicillin resistant *Staphylococcus aureus* enterocolitis. BMC Infect Dis 2015; 15: 265.

71 Karlsson FH, Ussery DW, Nielsen J, Nookaew I. A closer look at bacteroides: phylogenetic relationship and genomic implications of a life in the human gut. Microb Ecol 2011; 61: 473–485.

72 Bornbusch SL, Greene LK, McKenney EA, Volkoff SJ, Midani FS, Joseph G, Gerhard WA, Iloghalu U, Granek J, Gunsch CK. A comparative study of gut microbiomes in captive nocturnal strepsirrhines. Am J Primatol 2019; 81: e22986.

73 Greene LK, Clayton JB, Rothman RS, Semel BP, Semel MA, Gillespie TR, Wright PC, Drea CM. Local habitat, not phylogenetic relatedness, predicts gut microbiome structure within frugivorous and folivorous lemur lineages..

74 Macrina FL, Mays TD, Smith CJ, Welch RA. Non-plasmid associated transfer of antibiotic resistance in *Bacteroides*. J Antimicrob Chemother 1981; 8: 77–86.

75 Whittle G, Shoemaker NB, Salyers AA. The role of *Bacteroides* conjugative transposons in the dissemination of antibiotic resistance genes. Cell Mol Life Sci C 2002; 59: 2044–2054.

76 Privitera G, Dublanchet A, Sebald M. Transfer of multiple antibiotic resistance between subspecies of *Bacteroides fragilis*. J Infect Dis 1979; 139: 97–101.

77 Shoemaker NB, Vlamakis H, Hayes K, Salyers AA. Evidence for extensive resistance gene transfer among *Bacteroides spp*. and among *Bacteroides* and other genera in the human colon. Appl Environ Microbiol 2001; 67: 561–568.

78 Wexler HM. Bacteroides: the good, the bad, and the nitty-gritty. Clin Microbiol Rev 2007; 20: 593–621.

79 Veloo ACM, Baas WH, Haan FJ, Coco J, Rossen JW. Prevalence of antimicrobial resistance genes in *Bacteroides spp*. and *Prevotella spp*. Dutch clinical isolates. Clin Microbiol Infect 2019; 25: 1156–e9.

80 Stiefel U, Tima MA, Nerandzic MM. Metallo-β-lactamase-producing *Bacteroides* species can shield other members of the gut microbiota from antibiotics. Antimicrob Agents Chemother 2015; 59: 650–653.

81 Zhang A-N, Li L-G, Yin X, Dai CL, Groussin M, Poyet M, Topp E, Gillings MR, Hanage WP, Tiedje JM et al. Choosing your battles: which resistance genes warrant global action? BioRxiv 2019; : 784322.

82 Finnicum CT, Beck JJ, Dolan C V, Davis C, Willemsen G, Ehli EA, Boomsma DI, Davies GE, de Geus EJC et al. Cohabitation is associated with a greater resemblance in gut microbiota which can impact cardiometabolic and inflammatory risk. BMC Microbiol 2019; 19: 1–10.

83 Song SJ, Lauber C, Costello EK, Lozupone CA, Humphrey G, Berg-Lyons D, Caporaso JG, Knights D, Clemente JC, Nakielny S et al. Cohabiting family members share microbiota with one another and with their dogs. Elife 2013; 2: e00458.

84 Archie EA, Tung J. Social behavior and the microbiome. Curr Opin Behav Sci 2015; 6: 28–34.

85 Bennett PM. Plasmid encoded antibiotic resistance: acquisition and transfer of antibiotic resistance genes in bacteria. Br J Pharmacol 2008; 153.

86 Ventola CL. The antibiotic resistance crisis: part 2: management strategies and new agents. Pharm Ther 2015; 40: 344.

87 Van Puyvelde S, Deborggraeve S, Jacobs J. Why the antibiotic resistance crisis requires a One Health approach. Lancet Infect Dis 2018; 18: 132–134.

88 Bengtsson B, Greko C. Antibiotic resistance—consequences for animal health, welfare, and food production. Ups J Med Sci 2014; 119: 96–102.

89 Cabello FC. Heavy use of prophylactic antibiotics in aquaculture: a growing problem for human and animal health and for the environment. Environ Microbiol 2006; 8: 1137–1144.

90 Perry MR, McClean D, Simonet C, Woolhouse M, McNally L. Focusing on resistance to front-line drugs is the most effective way to combat the antimicrobial resistance crisis. bioRxiv 2018; : 498329.

91 Austvoll CT, Gallo V, Montag D. Health impact of the Anthropocene: the complex relationship between gut microbiota, epigenetics, and human health, using obesity as an example. Glob Heal Epidemiol Genomics 2020; 5.

92 Gillings MR, Paulsen IT. Microbiology of the Anthropocene. Anthropocene 2014; 5: 1–8.

93 Stearns SC. Evolutionary medicine: its scope, interest and potential. Proc R Soc B Biol Sci 2012; 279:4305–4321.

94 Trevathan WR. Evolutionary medicine. Annu Rev Anthropol 2007; 36.

95 Bornbusch SL, Grebe NM, Lunn S, Southworth CA, Dimac-Stohl K, Drea C. Stable and transient structural variation in lemur vaginal, labial and axillary microbiomes: patterns by species, body site, ovarian hormones and forest access. FEMS Microbiol Ecol 2020; 96: fiaa090.

96 Hernando-Amado S, Coque TM, Baquero F, Martínez JL. Defining and combating antibiotic resistance from One Health and Global Health perspectives. Nat Microbiol 2019; 4:1432–1442.

97 Trinh P, Zaneveld JR, Safranek S, Rabinowitz PM. One health relationships between human, animal, and environmental microbiomes: a mini-review. Front public Heal 2018; 6: 235.

98 Starling AP, Charpentier MJE, Fitzpatrick C, Scordato ES, Drea CM. Seasonality, sociality, and reproduction: long-term stressors of ring-tailed lemurs (*Lemur catta*). Horm Behav 2010; 57: 76–85.

99 Drea CM. Sex and seasonal differences in aggression and steroid secretion in *Lemur catta*: are socially dominant females hormonally ‘masculinized’? Horm Behav 2007; 51: 555–567.

100 Charles-Smith LE, Cowen P, Schopler R. Environmental and physiological factors contributing to outbreaks of *Cryptosporidium* in Coquerel’s Sifaka (*Propithecus coquereli*) at the Duke Lemur Center: 1999–2007. J Zoo Wildl Med 2010; 41: 438–444.

101 da Silva AJ, Cacciò S, Williams C, Won KY, Nace EK, Whittier C, Pieniazek NJ, Eberhard ML. Molecular and morphologic characterization of a *Cryptosporidium* genotype identified in lemurs. Vet Parasitol 2003; 111: 297–307.

102 Caporaso JG, Lauber CL, Walters WA, Berg-Lyons D, Huntley J, Fierer N, Owens SM, Betley J, Fraser L, Bauer M et al. Ultra-high-throughput microbial community analysis on the Illumina HiSeq and MiSeq platforms. ISME J 2012; 6: 1621.

103 Hall M, Beiko RG. 16S rRNA gene analysis with QIIME2. In: Microbiome Analysis. Springer, 2018, pp 113–129.

104 Callahan BJ, McMurdie PJ, Rosen MJ, Han AW, Johnson AJA, Holmes SP. DADA2: high-resolution sample inference from Illumina amplicon data. Nat Methods 2016; 13: 581.

105 Quast C, Pruesse E, Yilmaz P, Gerken J, Schweer T, Yarza P, Peplies J, Glöckner FO. The SILVA ribosomal RNA gene database project: improved data processing and web-based tools. Nucleic Acids Res 2012; 41: D590–D596.

106 Yarza P, Yilmaz P, Pruesse E, Glöckner FO, Ludwig W, Schleifer K-H, Whitman WB, Euzéby J, Amann R, Rosselló-Móra R. Uniting the classification of cultured and uncultured bacteria and archaea using 16S rRNA gene sequences. Nat Rev Microbiol 2014; 12: 635.

107 Hasan NA, Young BA, Minard-Smith AT, Saeed K, Li H, Heizer EM, McMillan NJ, Isom R, Abdullah AS, Bornman DM et al. Microbial community profiling of human saliva using shotgun metagenomic sequencing. PLoS One 2014; 9: e97699.

108 Yan Q, Wi YM, Thoendel MJ, Raval YS, Greenwood-Quaintance KE, Abdel MP, Jeraldo PR, Chia N, Patel R et al. Evaluation of the CosmosID bioinformatics platform for prosthetic joint-associated sonicate fluid shotgun metagenomic data analysis. J Clin Microbiol 2019; 57.

109 Chekabab SM, Lawrence JR, Alvarado A, Predicala B, Korber DR. A health metadata-based management approach for comparative analysis of high-throughput genetic sequences for quantifying antimicrobial resistance reduction in Canadian hog barns. Comput Struct Biotechnol J 2020.

110 Feehan A, Garcia-Diaz J. Bacterial, gut microbiome-modifying therapies to defend against multidrug resistant organisms. Microorganisms 2020; 8: 166.

111 Pedersen EJ, Miller DL, Simpson GL, Ross N. Hierarchical generalized additive models in ecology: an introduction with mgcv. PeerJ 2019; 7: e6876.

112 Gloor GB, Macklaim JM, Fernandes AD. Displaying variation in large datasets: plotting a visual summary of effect sizes. J Comput Graph Stat 2016; 25: 971–979.

113 Fernandes AD, Reid JN, Macklaim JM, McMurrough TA, Edgell DR, Gloor GB. Unifying the analysis of high-throughput sequencing datasets: characterizing RNA-seq, 16S rRNA gene sequencing and selective growth experiments by compositional data analysis. Microbiome 2014; 2: 15.

114 Gloor GB, Macklaim JM, Pawlowsky-Glahn V, Egozcue JJ. Microbiome datasets are compositional: and this is not optional. Front Microbiol 2017; 8: 2224.

