## Supplementary Material 1 for "Longitudinal effects of antibiotics and fecal transplant on lemur gut microbiota structure, associations, and resistomes"

**Supplementary Material 1** - Hierarchical Generalized Additive Models (HGAMs)

All models are structured following Pedersen *et al*., 2019 and we report model syntax for use in the R (ver. 4.0.2) via the gam(?) function in package {mgcv}. Models were run on data spanning the entire experiment and on subsets of data spanning the treatment period and/or recovery period.

Model for variation in alpha and beta diversity:

Diversity_metric ~ Experimental_group * Year + s(Day, by = Experimental_group) + s(Animal, by = Year, bs = "re") + s(Experimental_group, bs = "re"), method = “REML”

Model for variation in abundance of ABR genes:

ABR_genes ~ Experimetnal_group*Time_point + Previous_ABX_course + s(Animal, bs = "re") + s(Experimetnal_group, bs = "re") method = “REML”

Pedersen EJ, Miller DL, Simpson GL, Ross N. Hierarchical generalized additive models in ecology: an introduction with mgcv. *PeerJ* 2019; **7**: e6876.
