## Supplementary Tables 1 & 2 for "Longitudinal effects of antibiotics and fecal transplant on lemur gut microbiota structure, associations, and resistomes"

Table S1. Pairwise bacterial associations with >-0.5 or <0.5 mean correlation values across the treatment and recovery periods.

| **Pairwise comparisons – During ABX treatment and recovery periods** | | **mean correlation values** | | |
|---|---|---|---|---|
| **taxon 1** | **taxon 2** | **ABX** | **ABXFT** | **CON** |
| genus Bacteroides | genus Parabacteroides | 0.798 | 0.785 | - |
| genus Ruminococcaceae NK4A214 group | genus Ruminococcus 1 | 0.787 | - | - |
| genus Christensenellaceae R-7 group | genus Ruminococcaceae NK4A214 group | 0.665 | 0.766 | - |
| genus Ruminococcaceae NK4A214 group | family Erysipelotrichaceae | 0.618 | - | - |
| genus Blautia | genus Ruminococcaceae UCG-004 | 0.603 | - | - |
| genus Ruminococcaceae UCG-008 | genus Phascolarctobacterium | 0.597 | - | - |
| genus Roseburia | genus Ruminococcus 1 | 0.592 | - | - |
| genus Bacteroides | genus Barnesiella | 0.589 | 0.636 | - |
| genus Ruminococcus 1 | family Erysipelotrichaceae | 0.575 | - | - |
| genus Prevotellaceae NK3B31 group | genus Lachnospiraceae NK4A136 group | 0.574 | - | - |
| genus Cerasicoccus | order WCHB1-41 | 0.570 | 0.514 | 0.631 |
| genus Faecalibacterium | genus Ruminococcaceae NK4A214 group | 0.565 | - | - |
| genus Faecalibacterium | genus Ruminococcus 1 | 0.565 | - | - |
| genus Lachnospiraceae NK4A136 group | genus Ruminococcus 1 | 0.560 | - | - |
| genus Barnesiella | genus Desulfovibrio | 0.555 | - | - |
| genus Treponema 2 | family Bacteroidales RF16 group | 0.555 | - | - |
| genus Bacteroides | family Christensenellaceae | 0.551 | - | - |
| genus Bacteroides | family Clostridiales vadinBB60 group | 0.542 | - | - |
| genus Parabacteroides | genus Bilophila | 0.538 | 0.668 | - |
| genus Prevotellaceae NK3B31 group | genus Ruminococcaceae UCG-005 | 0.537 | - | - |
| genus Barnesiella | genus Parabacteroides | 0.531 | 0.607 | - |
| genus Sphaerochaeta | family Clostridiales vadinBB60 group | 0.524 | - | - |
| genus Parabacteroides | family Christensenellaceae | 0.520 | - | - |
| genus Christensenellaceae R-7 group | genus Ruminococcus 1 | 0.516 | - | - |
| genus Bacteroides | genus Bilophila | 0.515 | 0.645 | - |
| genus Bacteroides | genus UBA1819 | 0.514 | 0.620 | - |
| genus Angelakisella | genus Ruminiclostridium 5 | 0.514 | - | - |
| genus Roseburia | genus Ruminococcaceae NK4A214 group | 0.507 | - | - |
| genus Roseburia | genus Faecalibacterium | 0.501 | - | - |
| genus Ruminococcaceae UCG-008 | genus Other | -0.514 | - | - |
| genus Bacteroides | genus [Eubacterium] coprostanoligenes group | -0.534 | -0.557 | - |
| genus Bacteroides | genus Ruminococcaceae NK4A214 group | -0.540 | - | - |
| genus Bacteroides | genus Ruminococcus 1 | -0.542 | - | - |
| genus Parabacteroides | genus [Eubacterium] coprostanoligenes group | -0.551 | -0.527 | - |
| genus Bacteroides | genus Lachnospiraceae NK4A136 group | -0.555 | - | - |
| genus Ruminococcaceae NK4A214 group | genus Ruminococcaceae UCG-005 | - | 0.615 | - |
| genus Ruminococcaceae UCG-014 | family Lachnospiraceae | - | 0.544 | - |
| genus Bacteroides | genus Anaeroplasma | - | 0.540 | - |
| genus Parabacteroides | genus Coprococcus 1 | - | 0.529 | - |
| genus Barnesiella | genus Bilophila | - | 0.527 | - |
| genus Parabacteroides | genus UBA1819 | - | 0.519 | - |
| genus Candidatus Saccharimonas | family Lachnospiraceae | - | 0.511 | - |
| genus UBA1819 | genus Bilophila | - | 0.502 | - |
| genus Parabacteroides | genus Ruminococcaceae UCG-005 | - | -0.518 | - |
| genus Rikenellaceae RC9 gut group | genus Bilophila | - | -0.520 | - |
| genus Parabacteroides | genus Ruminococcaceae NK4A214 group | - | -0.530 | - |
| genus Prevotella 9 | genus Other | - | -0.533 | - |
| genus Rikenellaceae RC9 gut group | genus Parabacteroides | - | -0.548 | - |
| genus Bacteroides | genus Ruminococcaceae UCG-005 | - | -0.555 | - |

| | | | | |
|---|---|---|---|---|
| genus Bacteroides | genus Rikenellaceae RC9 gut group | - | -0.561 | - |
| genus Bacteroides | family Lachnospiraceae | - | -0.569 | - |
| genus Ruminococcaceae UCG-014 | genus Bilophila | - | -0.570 | - |
| genus Ruminococcaceae UCG-014 | genus UBA1819 | - | -0.587 | - |
| genus Parabacteroides | family Lachnospiraceae | - | -0.606 | - |
| genus Parabacteroides | genus Ruminococcaceae UCG-014 | - | -0.624 | - |
| genus Bacteroides | genus Ruminococcaceae UCG-014 | - | -0.642 | - |

Table S2. Pairwise bacterial associations with >-0.5 or  <0.5 mean correlation values across the recovery periods.

| **Pairwise comparisons - Recovery period** | | **mean correlation values** | | |
|---|---|---|---|---|
| **taxon 1** | **taxon 2** | **ABX** | **ABXFT** | **CON** |
| genus Bacteroides | genus Parabacteroides | 0.726 | 0.713 | - |
| genus Prevotellaceae NK3B31 group | genus Ruminococcaceae UCG-005 | 0.651 | - | - |
| genus Succinivibrio | genus Treponema 2 | 0.620 | - | - |
| genus Bacteroides | genus Angelakisella | 0.617 | - | - |
| genus Ruminiclostridium 5 | genus Bilophila | 0.606 | - | - |
| family Muribaculaceae | genus Prevotellaceae NK3B31 group | 0.552 | - | - |
| family Bacteroidales RF16 group | genus Treponema 2 | 0.543 | - | - |
| genus Blautia | genus Ruminococcaceae UCG-004 | 0.536 | - | - |
| genus Bacteroides | genus Anaeroplasma | 0.531 | - | - |
| genus Rikenellaceae RC9 gut group | genus [Eubacterium] coprostanoligenes group | 0.529 | - | - |
| genus Treponema 2 | genus Cerasicoccus | 0.528 | - | - |
| genus Parabacteroides | genus Angelakisella | 0.521 | - | - |
| genus Oscillospira | genus Ruminiclostridium 5 | 0.518 | - | - |
| genus Lachnoclostridium | genus Oribacterium | 0.513 | - | - |
| genus Christensenellaceae R-7 group | genus Ruminococcaceae NK4A214 group | 0.506 | 0.518 | - |
| genus Parabacteroides | genus Oscillospira | 0.502 | - | - |
| genus Blautia | genus GCA-900066575 | 0.501 | - | - |
| genus p-1088-a5 gut group | genus Treponema 2 | 0.500 | - | - |
| genus Prevotella 9 | genus Ruminiclostridium 5 | -0.521 | - | - |
| genus Parabacteroides | genus Sarcina | -0.536 | - | - |
| genus Bacteroides | genus Ruminococcaceae UCG-008 | - | 0.648 | - |
| order WCHB1-41 | genus Cerasicoccus | - | 0.629 | 0.568 |
| genus Butyricimonas | genus Prevotella 1 | - | 0.605 | - |
| genus Bacteroides | genus Roseburia | - | 0.595 | - |
| genus Oscillospira | genus Ruminococcaceae NK4A214 group | - | 0.592 | - |
| genus Parabacteroides | genus Coprococcus 1 | - | 0.576 | - |
| genus Ruminococcaceae NK4A214 group | genus Ruminococcaceae UCG-005 | - | 0.575 | - |
| genus Rikenellaceae RC9 gut group | genus Treponema 2 | - | 0.572 | - |
| genus Oscillospira | genus Ruminococcaceae UCG-005 | - | 0.566 | - |
| genus Parabacteroides | genus Bilophila | - | 0.561 | - |
| genus Bacteroides | genus Bilophila | - | 0.547 | - |
| genus Cerasicoccus | genus [Eubacterium] coprostanoligenes group | - | 0.534 | - |
| genus Parabacteroides | genus Lachnospiraceae UCG-001 | - | 0.533 | - |
| family Muribaculaceae | genus Ruminococcaceae UCG-013 | - | 0.514 | - |
| genus Bacteroides | genus Barnesiella | - | 0.513 | - |
| genus Prevotellaceae UCG-003 | genus Blautia | - | 0.513 | - |
| genus Roseburia | genus Ruminococcaceae UCG-008 | - | 0.512 | - |
| genus Bacteroides | genus Lachnospiraceae UCG-001 | - | 0.512 | - |
| genus Bacteroides | genus UBA1819 | - | 0.512 | - |
| family Muribaculaceae | genus Lachnospiraceae NK4A136 group | - | 0.511 | - |

| | | | | |
|---|---|---|---|---|
| genus Lachnoclostridium | genus Ruminococcaceae UCG-008 | - | 0.506 | - |
| genus Ruminococcaceae UCG-014 | genus Bilophila | - | -0.509 | - |
| genus Bacteroides | family Lachnospiraceae | - | -0.515 | - |
| genus Bacteroides | genus [Eubacterium] coprostanoligenes group | - | -0.520 | - |
| genus Parabacteroides | family Lachnospiraceae | - | -0.534 | - |
| genus Lachnospiraceae UCG-001 | genus Ruminococcaceae UCG-014 | - | -0.542 | - |
| genus Parabacteroides | genus Ruminococcaceae UCG-014 | - | -0.551 | - |
| genus Bacteroides | genus Ruminococcaceae UCG-014 | - | -0.572 | - |
| genus Ruminococcaceae UCG-014 | genus UBA1819 | - | -0.597 | - |
| genus Cerasicoccus | order Rhodospirillales | - | - | 0.532 |
